# Validation of a male-specific DNA marker confirms XX/XY-type sex determination in African catfish (*Clarias gariepinus*)

**DOI:** 10.1101/2022.10.12.511891

**Authors:** Réka Enikő Balogh, Balázs Csorbai, Csaba Guti, Szilvia Keszte, Béla Urbányi, László Orbán, Balázs Kovács

## Abstract

African catfish (*Clarias gariepinus*) is a promising food fish species with significant potential and growing mass of production in freshwater aquaculture. Male African catfish possess improved production characteristics over females, therefore the use of monosex populations could be advantageous for aquaculture production. However, our knowledge about the sex determination mechanism of this species is still limited and controversial. A previously isolated male-specific DNA marker (CgaY1) was validated using offspring groups from targeted crosses (n=630) and it was found to predict the sex of 608 individuals correctly (96.43% accuracy). Using the proportion of recombinants, we estimated the average genetic distance between the potential sex determination locus and the sex-specific marker to be 2.69 cM. As an earlier study suggested that both XX/XY and ZZ/ZW systems coexist in this species, we tested the applicability of their putative ‘moderately sex-linked loci’ and found that no sex-specific amplification could be detected for any of them. In addition, temperature-induced masculinization suggested by others was also tested, but no such effect was detected in our stocks when the published parameters were used for heat treatment. Altogether, our results support an exclusive XX/XY sex determination system in our African catfish stock and indicate a good potential for the future use of this male-specific DNA marker in research and commercial production.

## 1. Introduction

The estimated number of fish species on Earth is well over 32,000 (Nelson, 2016) and they inhabit habitats with wide-ranging conditions. They show a very diverse range of phenotypes and possess heterogeneous and highly flexible sex determination (SD) mechanisms, including systems based on genetic and environmental SD, as well as sex chromosomal or polygenic SD systems (Paul-Prasanth et al., 2011; Penman and Piferrer, 2008). Nearly 400 species of the finfishes are considered to be commercially significant according to the UN Food and Agriculture Organization (FAO, 2018) and many of them display substantial sexual dimorphism in commercially important traits, such as growth rate (Beardmore et al., 2001; Kocour et al., 2003; Mei and Gui, 2018; Zhou et al., 2021), fillet quality (Manor et al., 2015), yield (Bosworth et al., 2001) or mortality rate (Su et al., 2013). Therefore, the use of monosex production for these species can be desirable.

However, our knowledge about the genetic background of sex determination in fishes is still limited, even for species with the less elusive monofactorial genetic SD system. Heretofore, only a few master SD genes have been published for teleosts: *dmy* for *Oryzias latipes* and *O. curvinotus* (Matsuda et al., 2002; Volff et al., 2007), *gsdf*^Y^ in *O. luzonensis* (Myosho et al., 2012), *amhY* in *Odontesthes hatcheri* (Hattori et al., 2012), *sdY* in *Oncorhynchus mykiss* (Yano et al., 2012), *pfpdz1* for *Pelteobagrus fulvidraco* (Dan et al., 2018), *amh* for *Esox lucius* (Pan et al., 2019), and lastly, a missense single-nucleotide polymorphism (SNP) in *amhr2* in *Takifugu rubripes* (Kamiya et al., 2012). Some of these SD genes have been shown to be conserved in several related species; *sdY* turned out to be the primary switch in an additional ten salmonid species as well (Yano et al., 2013). Most of these ‘master switches’ are paralogs created by the teleost-specific genome duplication (Heule et al., 2014; Schartl, 2004; Volff et al., 2007) and they can encode ‘modified’ transcription factors (Matsuda et al., 2002; Volff et al., 2007), secretory proteins (Hattori et al., 2012; Myosho et al., 2012), and hormone receptors (Kamiya et al., 2012). Apparently - in contrast with the highly conserved male and female heterogamety in mammals and birds - a wide variety of signals can control SD and early gonad differentiation in fish [for reviews see: (Yano et al., 2012; Dan et al., 2018; Pan et al., 2019)]. Therefore, our chances for identifying a universal SD mechanism in teleosts are dim and in the absence of known SD genes the potential use of sex-specific markers gains higher importance. Despite a wide range of potential applications in both commercial production and research, routine use of such markers in either of these areas is very limited.

Sex-specific DNA markers, including Amplified Fragment Length Polymorphism [AFLP; (Mueller and Wolfenbarger, 1999)], Random Amplified Polymorphic DNA [RAPD; (Hadrys et al., 1992)], Restriction Fragment Length Polymorphism [RFLP; (Tanksley et al., 1989)], Sequence Characterized Amplified Region [SCAR; (Kiran et al., 2010)] and Simple Sequence Repeat [SSR; (Powell et al., 1996)] markers have been published for a number of fish, including eleven catfish species (see **Table 1** for details). For instance, in yellow catfish (*Pelteobagrus fulvidraco*) two SCAR markers were successfully used to help the production of all-male populations, which have a great potential in aquaculture, as adult males can grow over twice the size that of females (Wang et al., 2009). In fathead minnow (*Pimephales promelas* a sex-linked AFLP marker was used for research purposes to detect sex-reversal caused by endocrine chemical treatment (Olmstead et al., 2011). In rainbow trout (*Oncorhynchus mykiss*), a SCAR marker was found to predict sex with 92-100% accuracy in specific strains (Felip et al., 2005). These markers may offer higher reliability in comparison to sexing based on phenotypic traits. In channel catfish (*Ictalurus punctatus*), sexing based on external characteristics corresponded to the internal anatomy in only 70% of the investigated specimens, whereas a sex-linked microsatellite marker (AUEST0678) offered 100% fidelity (Ninwichian et al., 2012).

**Table 1:** Sex-specific genomic DNA markers isolated from catfish (Siluriformes) genomes

| Species | Family | Type of isolated marker | Sex chrom. system | Number of markers | Conversion to locus-specific marker | Best efficiency | Validation (number of specimen) | Multi-generational testing | References |
| --- | --- | --- | --- | --- | --- | --- | --- | --- | --- |
| *Clarias gariepinus* | *Clariidae* | RAPD | XX/XY | 2 | Yes | 100%<br>95% | Yes (20-20) | No | (Kovács et al., 2001) |
| *Clarias gariepinus* | *Clariidae* | SCAR | XX/XY | 1* | Yes | 96.4% | Yes (630) | Yes | present study |
| *Clarias gariepinus* | *Clariidae* | PA & SNP | XX/XY and ZZ/ZW | 41 & 25 | No | No data | No | No | (Nguyen et al., 2021a) |
| *Clarias fuscus* | *Clariidae* | SNP | XX/XY | 5 | Yes | 100% | Yes (84) | No | (Lin et al., 2022) |
| *Clarias macrocephalus* | *Clariidae* | PA & SNP | XX/XY | 17 & 7 | Yes | 100% | Yes (30) | No | (Nguyen et al., 2021b) |
| *Pelteobagrus fulvidraco* | *Bagridae* | AFLP | XX/XY | 6 | Yes | 100% | Yes (577) | No | (Wang et al., 2009) |
| *Pelteobagrus fulvidraco* | *Bagridae* | AFLP | XX/XY | 3 | Yes | 100% | Yes (80) | No | (Dan et al., 2013) |
| *Pelteobagrus vachelli* | *Bagridae* | SNP | XX/XY | 12 | Yes | 100% | No | No | (Zhang et al., 2020) |
| *Pseudobagrus ussuriensis* | *Bagridae* | Microsat. | XX/XY | 1 | Yes | 100% | Yes (185) | No | (Zhu et al., 2018) |
| *Hemibagrus wyckioides* | *Bagridae* | 2b-RAD-tags/SNP | XX/XY | 67 | No | 44.2-85.7% | No | No | (Zhou et al., 2021) |
| *Leiocassis longirostris* | *Bagridae* | 2b-RAD-tags | XX/XY | 1 | Yes | 100% | Yes (230) | No | (Dai et al., 2022) |
| *Ictalurus punctatus* | *Ictaluridae* | Microsat. & SNP/SCAR | XX/XY | 25 | Yes | 100% | Yes (80) | No | (Ninwichian et al., 2012) |
| *Silurus lanzhouensis* | *Siluridae* | 2b-RAD-tags/SNP | XX/XY | 3 | Yes | 100% | Yes (76) | Yes* | (Wang et al., 2021) |
| *Silurus meridionalis* | *Siluridae* | k-mer | XX/XY | 8 | Yes | 100% | Yes (165) | No | (Zheng et al., 2020) |

\* Testing in two generations

African catfish (*Clarias gariepinus*) belongs to the Siluriformes order and the *Clariidae* family. It is an important cultured freshwater species with growing mass of production in numerous African, South American, and European countries (FAO, 2022; OECD, 2022). Rapid growth rate, efficient feed conversion, omnivoracity, and high resistance to diseases make this species highly suitable for intensive aquaculture. Sex-related differences exist in production characteristics: males exhibit higher growth rate, better feed utilization (Henken et al., 1987) and higher fillet yield (Oellermann and Hecht, 2000). Although rearing all-male populations could have significant economic implications, according to our knowledge only mixed populations exist at commercial fish farm.

Apart from being a notable commercial food fish species, African catfish is also widely used as a model organism in a number of toxicological, morphological and physiological studies (Hassan et al., 2018; Sayed, 2016; Verreth and Den Bieman, 1987). Despite the importance of this gonochoristic species, there was a debate earlier about its sex determination mechanism. Some of the researchers suggested a ZZ/ZW SD system (Ozouf-Costaz et al., 1990, Váradi et al., 1999) based on karyological analysis and the sex-ratio of gynogenetic offspring, whereas others proposed the presence of an XX/XY-type one (Liu and Yao, 1995, Galbusera et al., 2000) based on the sex-ratio of offspring after gynogenesis and self-fertilization. Recently, it was suggested that both XX/XY and ZZ/ZW systems coexist in Asian populations of African catfish based on a study where 41 ‘moderately male-linked’ and 25 ‘moderately female-linked’ loci were isolated with Diversity Arrays Technology using 30 specimens (Nguyen et al., 2021a). Earlier, we isolated a male-specific DNA marker (CgaY1) from African catfish (Kovács et al., 2001), that could be used for molecular sexing from as early as zygote stage. Our data indicated that the species or at least the stock analyzed in those experiments had an XX/XY sex chromosomal SD system.

Moreover, in addition to genetic sex determination (GSD), temperature-based sex reversal was also suggested in African catfish (Santi et al., 2016; Santi et al., 2017) with masculinizing effect of high temperature, though this hypothesis has not been confirmed with molecular techniques yet. African catfish larvae were exposed to high temperature (36°C) for three days at different developmental stages and they suggested the most thermosensitive period to be 6-8 days post-hatching (dph) (Santi et al., 2016). They found skewed sex-ratios (90-100% of males) in 10 of 19 families examined. However, the authors also observed high variability of sex ratios between families, therefore they proposed a potential parental effect in the thermosensitive process.

In the present study, we have set out to validate a previously isolated male-specific SCAR marker (CgaY1; GenBank accession number: AF332597) (Kovács et al., 2001) on African catfish offspring produced by targeted crosses. Moreover, the possibility for temperature-dependent sex reversal suggested by others for this species (Santi et al., 2016) was also analyzed with molecular techniques. Besides, the applicability of four recently published putative ‘moderately sex-associated markers’ (Nguyen et al., 2021a) was tested on our African catfish stock to investigate the SD mechanism of this species.

## 2. Materials and methods

### 2.1 Origin of the brooders

Seven specimens of P0 African catfish brooders were obtained from four different Hungarian broodstocks: one from Jászkiséri Halas Ltd. (Jászkisér), one from V-95 Ltd. (Nagyatád), and five from Szarvas Fish Ltd. (three from Tuka and two from Szarvas, respectively). The stock at Szarvas (GPS: 46°52′0.01″ N 20°33′0.00″ E) was established in the 1980s from specimens originating from the Netherlands. Some brooders were added unsystematically during the following years. The broodstock at Tuka (GPS: 47°44′31.3″ N 21°02′06.2″ E) was set up from a stock originating from Denmark in 1988, which was expanded later with specimens from Szarvas (Ferenc Radics, personal communication). The stock in Jászkisér (GPS: 47°28′51.0″ N 20°15′22.3″ E) was founded with brooders purchased from Szarvas (Gyula Borbély, personal communication). The stock at Nagyatád (GPS: 46°12′34.1″ N 17°22′25.5″ E) was set up with brooders purchased from several Hungarian fish breeders (Attila Boros, personal communication).

### 2.2 Fish rearing and genetic crosses for validation of the CgaY1 marker

Two experiments were performed at the Szent István Campus with the permission PEI/001/1719-2/2015 for animal research. In the first one (**Figure 1A**; Experiment #1), 179 F1 siblings from a pairwise cross were sexed molecularly using the CgaY1 DNA marker 3-5 weeks after propagation (see Section 2.5 for a detailed protocol). Individuals were sorted according to their ‘molecular sex’ into a male group and a female group, respectively, and the groups were subsequently reared separately in 0.6 m^3^ tanks, at 28°C, with 16L/8D light cycle and with complete water change once a week. Their phenotypic sex was determined five months post-hatching (mph) with aceto-carmine squash method (see Section 2.4 for a detailed protocol).

**Figure 1:**
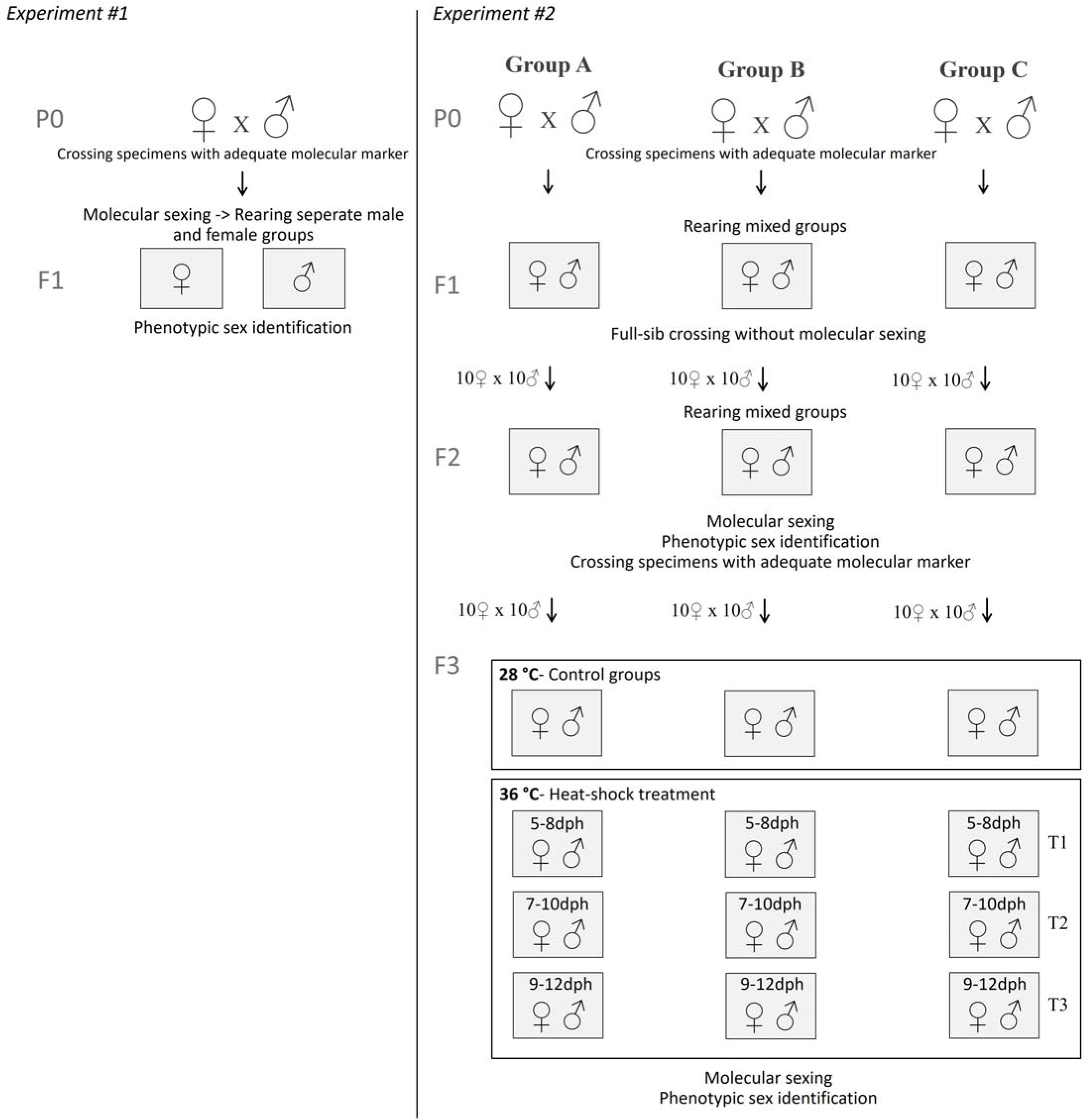
Schematic representation of the genetic crossing and sampling methods used to produce the targeted crosses for the validation of the CgaY1 marker in African catfish. The F1 generation was analyzed in Experiment #1, whereas the F2 generation and control groups of F3 generation in Experiment #2. For the investigation of the heat-induced sex reversal, individuals of the F3 generation (Experiment #2) were used.

In the second experiment (**Figure 1B**; Experiment #2), all brooders were tested for the presence of the male-specific sex marker (CgaY1) by PCR prior to the crosses. Three separate pairwise crosses were made using five specimens as brooders (one of them was used in two different crosses), then the resulting F1 offspring were grown in three different tanks to maturation without subjecting them to molecular sexing. The F2 generation was produced by pairwise full-sib crossing of randomly selected specimens from the F1 generation (ten males and ten females from each cross) resulting in a total of 30 families. F2 specimens were then reared in mixed-sex groups in three different tanks (one tank for each cross) until maturation and they were sexed by using the molecular marker. Afterwards, their phenotypic sex was identified by visual observation of gonadal morphology following dissection and ten females and ten males were selected from all three F2 groups with the corresponding molecular marker to produce ten F3 families in each group. The combined F3 offspring population from each of these groups was reared in mixed-sex groups in three different tanks (one for each group) until three months post-hatching (mph) and then, their phenotypic sex was determined by (i) aceto-carmine squash method; and their genotypic sex by (ii) sexing with the CgaY1 molecular marker.

All F2 families were raised in a recirculating aquaculture system in 1 m^3^ tanks, at 25°C, with 16L/8D light cycle and with a replacement water flow rate of 80 liters/hour. F3 families were raised in recirculated tanks of 12 liter volume, with 1000 juvenile fish in each tank, at 28°C, with 16L/8D light cycle and with a replacement water flow rate of 80 liters/hour. All in all, three independent F2 batches (85, 86, and 89 specimens, respectively) and three independent F3 batches (50, 48, 93 specimens, respectively) were analyzed in the second experiment to validate the CgaY1 male-specific marker.

### 2.3 Early heat treatment

In order to study the heat-induced sex reversal in African catfish recommended earlier (Santi et al., 2016), juveniles from the F3 generation were subjected to heat shock treatment (36±1°C) for 72 hours at three different developmental stages. Three batches of 1000 larvae each were created from the three F3 crosses (Cross F3A, F3B and F3C) consisting of a total of 30 families (10 families from each cross, see **Figure 1C**). Then three groups of ca. 1000 individuals each for each cross were exposed to increased temperature (36±1°C) at different developmental stages: T1 – 5-8 days post-hatching (dph), T2 – 7-10 dph and T3 – 9-12 dph. Experimental temperature (36±1°C) was reached in a water bath within 3 hours and the temperature of the individual tanks was monitored by RC-4 temperature data logger (Elitech). Water was cooled down to 28°C within 24 hours after each treatment, then the fish were raised in nine separate tanks (12 liters each) in recirculating aquaculture system (RAS), at 28°C, with 16L/8D light cycle and with a replacement water flow rate of 80 liters/hour. Fish were grown until three mph and the phenotypic sex of all individuals was determined by aceto-carmine squash method. A total of 123 phenotypic males (24, 19 and 80 fish from the three crosses, respectively) were randomly selected from these heat-treated groups and sexed by the CgaY1 molecular marker.

### 2.4 Determination of phenotypic sex

In the F1 and F3 groups, phenotypic sex was determined 3-5 weeks and 3-4 months after hatching, respectively by a modified aceto-carmine squash method (Guerrero and Shelton, 1974) using indigo-carmine dye instead of carmine in order to obtain a better contrast. Tissues were investigated under a stereomicroscope (Leica M205FA) with 40-100X magnification (see **Figure 2** for typical examples). In F2 families, mature individuals were sacrificed by overdosing with 2-phenoxyethanol (Sigma-Aldrich), then the two sexes were identified by examining gonadal morphology following dissection.

### 2.5 Molecular sexing using the CgaY1 marker

Fin clip tissues were collected for genetic analyses and stored in 98% ethanol at-20°C until use. DNA was isolated with the basic phenol-chloroform method (Blin and Stafford, 1976) from the F1 individuals and qualification and quantification of DNA were performed with NanoDrop One spectrophotometer (Thermo Scientific). On the F2 and F3 samples a rapid Chelex100-based procedure (Walsh et al., 1992) was applied. A duplex PCR reaction was performed amplifying the CgaY1 marker (approx. 1.5 kb) and a K1 control fragment (345 bp) for males and only the control (K1) fragment for females as described earlier (Kovács et al., 2001). One newly designed primer sequence was utilized: Y1-6F (5’-CCGTAGCCTGCCACTCTGC-3’) and the others were obtained from a previous study (Kovács, 2004):, Y1-1R (5’- GAACTGAACCCCACATTTTGTC-3’), K1-1F (5’-AGTACATTGAGGACGAGGACGC-3’), K1-1R (5’- CATTGTAACAAGAGGAGCCCAC- 3’). The optimal master mix for this duplex reaction was: 0.9X Promega PCR buffer, 200-200 μM of each dNTP, 500-500 nM Y1-5F and Y1-1R primers, 30-30 nM of K1-1F and K1-1R primers, 2mM MgCl2, 1U Taq DNA polymerase (Promega) and 20 ng genomic DNA. Cycling conditions were as follows: a pre-amplification denaturation at 94°C for 2 mins, followed by 30 cycles of 94°C for 30 s, 62.5°C for 1 min and 72°C for 3 mins, then a final extension of 72°C for 5 mins by using ProFlex (Applied Biosystems) PCR machine. Five μl of the resulting products were separated by agarose gel electrophoresis using 1.5% agarose (SeaKem LE, 1x TBE buffer (108g Tris, 55g boric acid, 9.3g EDTA per liter), 0.5μg/ml ethidium bromide, 5X Loading Dye (Thermo Scientific) and 5X GeneRuler DNA Ladder Mix (Thermo Scientific) with Consort Electrophoresis Power Supply E835 (Sigma Aldrich).

### 2.6 Investigation of putative sex-associated markers

The applicability of recently published ‘moderately sex-associated markers’ (Nguyen et al., 2021a) was tested on eight male and eight female African catfish specimens from the F2 generation of the targeted crosses. Fin clip tissues were collected and stored in 98% ethanol at-20°C until DNA isolation, which was carried out with E.Z.N.A Tissue DNA kit (Omega Bio-tek, Norcross, GA, USA). Qualification and quantification of DNA were performed with NanoDrop One spectrophotometer (Thermo Scientific) and by running 2ng DNA on agarose gel electrophoresis using 1% agarose, 1x TBE buffer and 0.5μg/ml ethidium bromide. We have analyzed three ‘moderately male-linked loci’: *dtna, add3* and *gucd1*, and two ‘moderately female-linked loci’: *dctn4* and *pcdh2ab3* described earlier (Nguyen et al., 2021a) (see **Supplementary Table S1** for additional information about the markers). The trimmed product sequences were searched using BLAST software with nucleotide algorithm on our unpublished African catfish genome. Hits were considered significant with E-value < 1. For four sequences a single significant alignment each was detected, whereas for *gucd1* no significant hit was found. Thirty primer pairs were designed for the flanking regions (within 100 bp in both directions) of each marker by Primer-BLAST software and the most specific primer pairs (**Supplementary Table S1.**) displaying significant hit only for the corresponding product were chosen for further analysis. The optimal master mix for this reaction was: 10X Taq buffer with KCl (Thermo Scientific), 200-200 μM of each dNTP, 252nM of F and R primers, 1.5mM MgCl_2_, 1U Taq DNA polymerase (Promega) and 100 ng genomic DNA. The optimal cycling conditions were the following: a pre-amplification denaturation at 95°C for 2 mins, followed by 30 cycles of 94°C for 30 s, 55°C for 20 s and 72°C for 1 min, then a final extension of 72°C for 10 mins by using ProFlex (Applied Biosystems) PCR machine. Five μl of the resulting PCR products were separated by agarose gel electrophoresis as described above.

### 2.7 Data analyses

The phenotypic sex was compared to the result of molecular sexing obtained by the CgaY1 sex-specific marker following PCR amplification. The average genetic distance between the potential sex determining locus and the male-specific CgaY1 marker was calculated from the recombination rates detected in the F2 generation. Deviations from 1:1 sex ratio were tested in every generation by one-proportion Z-test with continuity correction, and with 5% significance level using R software 3.5. Percentages of phenotypically male specimens not displaying the male-specific marker in the heat treated and control groups of F3 generation were compared by two-proportion Z-test for equal percentages with 5% significance level using R software 3.5.

## 3. Results and Discussion

This study had three aims: (1) validate our previously published Y-specific marker on African catfish families (Kovács et al., 2001); (2) test whether temperature-induced sex-reversal described earlier for this species (Santi et al., 2016) can be detected with this marker; and (3) test on our stocks some of the putative ‘moderately sex-linked markers’ isolated by others (Nguyen et al., 2021a).

Here, we report about the validation of the CgaY1 male-specific DNA marker that was performed on a total of 630 individuals originating from a large number of crosses spanning three generations. By comparing their phenotype with the data from molecular sexing, the latter was found to predict phenotypic sex with 96.43% fidelity, which is proportional to the results found with similar markers in other species (Felip et al., 2005; Ninwichian et al., 2012; Wang et al., 2009). We found a difference between the phenotypic and the genotypic sex in only 22 out of 630 individuals (3.57%). The Y-specific DNA marker could not be amplified from 10 phenotypic males, whereas templates of 12 phenotypic females produced the male-specific PCR fragment. All these 20 cases were confirmed by a second PCR reaction.

Neither genotypic nor phenotypic sex ratios differed significantly (p > 0.05) from 1:1 ratio in any of the groups examined (**Table 2**). Based on the data available from all from siluroids (**Table 1**), CgaY1 is the first molecular marker in *C. gariepinus* that has been validated and tested in multiple generations, and the second such marker from any catfish species behind the one reported by Wang and colleagues recently from Lanzhou catfish (Wang et al., 2021). In general, XX/XY sex determination system seems to be characteristic in the Siluriformes order based on data obtained with the previously isolated sex-related markers (**Table 1**).

**Table 2:** Segregation data of the CgaY1 male-specific genomic DNA marker and the sex ratios

| Batches analyzed | n | Recombination events | Fidelity (%) | CgaY1-positive males | CgaY1-positive females | Genotypic sex ratio (male %) | Phenotypic sex ratio (male %) |
| --- | --- | --- | --- | --- | --- | --- | --- |
| F1 | 179 | 6 | 96.65 | 82/87 | 1/92 | 57.7 | 58.8 |
| F2 Cross A | 85 | 1 | 98.82 | 49/50 | 0/35 | 39.5 | 37.6 |
| F2 Cross B | 86 | 4 | 95.35 | 31/32 | 3/54 | 51.7 | 57.5 |
| F2 Cross C | 89 | 2 | 97.75 | 46/48 | 0/41 | 49.7 | 48.6 |
| F3 Cross A | 50 | 2 | 96.00 | 29/29 | 2/21 | 58.0 | 62.0 |
| F3 Cross B | 48 | 2 | 95.83 | 28/28 | 2/20 | 58.3 | 62.5 |
| F3 Cross C | 93 | 5 | 94.62 | 45/46 | 4/47 | 52.0 | 58.0 |
| All | 630 | 22 | 96.43 | 310/320 | 12/310 | 52.4 | 55.0 |

Although the observed differences in the phenotypic and genotypic sex revealed by the CgaY1 male-specific marker can possibly result from environmental sex reversal, the probability of such event is relatively low, as all individuals were grown under the same environmental conditions. It is more likely, that recombination events occurred between the SD locus and the sex-specific marker in these individuals. In this case, the recombination frequencies between these two loci in the F2 generation were 1.18%, 4.65%, and 2.25%, respectively, and the estimated mean genetic distance between these two genomic positions is 2.69 cM. When F3 families were analyzed, they showed a slightly higher, but comparable average recombination frequency of 4.52%. Altogether, these results indicate strong linkage between the CgaY1 marker and the SD locus, and thereby offering opportunities for the applicability of this molecular sex marker in both production and research of African catfish. In aquaculture settings, the CgaY1 molecular sex marker offers a fast and cost-effective solution for molecular sexing during the early developmental stages, which could be used to validate monosex groups generated by hormonal treatments or genome manipulation. It could also have significant implications in research as well, for instance in genetic mapping or in ecotoxicological or developmental studies.

A marker closely associated with the SD enables the detection of temperature-dependent sex reversal, therefore, it is suitable for the verification of this phenomenon in this species. The possibility of temperature-induced masculinization reported earlier (Santi et al., 2016) was tested on the F3 generation of our targeted crosses at three different developmental stages (5-8 dph, 7-10 dph, 9-12 dph). When 123 heat-treated, phenotypic males were genotyped with the CgaY1 molecular marker, four individuals (3.25%) failed to amplify the male-specific PCR fragment. This is slightly higher, but not significantly different from the deviation experienced in the control group of the F3 generation, where 1% of the examined 103 males did not amplify the marker. The phenotypic sex ratios did not differ significantly (p > 0.05) from the 1:1 ratio in either of the three heat-treated groups (44%, 53%, 44% of males, respectively). Altogether, our results do not support the idea that temperature-induced sex reversal took place in our African catfish stock with the given parameters. However, a strong parental effect has been suggested for the heat-induced sex reversal in many fish species displaying GSD with temperature effect (Baroiller et al., 2009; Shen and Wang, 2014), whereas temperature-sensitive and -insensitive populations have been observed in other species (Valenzuela and Lance, 2004). Besides, many morphological malformities and a relatively high mortality rate were experienced at 90 dph in the heat-treated groups (82±3%) compared to those of the control groups (51±5%), which is comparable to those found in previous studies (Santi et al., 2016; Santi et al., 2017). Consequently, the applicability of this heat-based masculinization method for all-male production seems to be limited to some stocks or families (Santi et al., 2016) and not applicable to the stocks used by us.

More and more assembled genome sequences are becoming available from teleost species, and they enable us to perform large-scale *in silico* analyses, including pan-genome surveys (Vernikos et al., 2015), marker development and cross-species applications (Luo et al., 2012; You et al., 2020), detection of sex determination mechanisms (Xu et al., 2020) and evolutionary studies (Christoffels et al., 2004). According to our knowledge, complete genomes from at least 22 species of the Siluriformes order have been sequenced and assembled so far [see e.g. (Chen et al., 2016; Gao et al., 2021; Li et al., 2018; Ozerov et al., 2020; Zhou et al., 2018)], allowing for identification of potential orthologues of genes with sex-related or sex-specific function. Our group performed the whole genome sequencing of a male African catfish specimen (B. Kovács, unpublished data), which is also expected to provide interesting possibilities for further studies along these lines.

Earlier investigations yielded contradictory results about the sex determination mechanism of African catfish supporting either XX/XY or ZZ/ZW systems. Recently, a report has been published about the analysis of sex chromosomal systems in this species (Nguyen et al., 2021a). The authors have used an approach called Diversity Arrays Technology (DArT) that is based on ‘high-throughput genome complexity reduction sequencing’ to search for sex-associated DNA markers in the genome of African catfish individuals obtained from the broodstock of Kasetsart University (Thailand; the origin of the stock was not indicated in the publication). Through the analysis of thirty phenotypically sexed individuals, they have identified 41 ‘moderately male-linked loci’ and 25 ‘moderately female-linked loci’ and suggested, that both XX/XY and ZZ/ZW sex-determination mechanisms co-exist in the African catfish stocks analyzed. In the present study, we tried to validate in our stocks three of their ‘moderately male-linked loci’: *gucd1, add3* and *dtna*, as well as two ‘moderately female-linked loci’: *dctn4* and *pcdh2ab3* (for gene names and primer sequences see **Supplementary Table S2**). As one of these putative markers, *gucd1*, could not be found in our *C. gariepinus* genome (unpublished data), the applicability of the remaining four putative markers for molecular sexing was checked on eight mature female and eight mature male specimens. All four PCR reactions yielded a single product of equal size each from both sexes (**Supplementary Figure S2**). We concluded that these ‘moderately sex-linked’ markers cannot be used for sexing in our African catfish stock, and consequently, these results do not support the presence of a ZZ/ZW sex-determination system in our African catfish stock.

In summary, the CgaY1 male-specific molecular marker provides a reliable, fast, and cost-effective way for testing the sex differentiation process and its outcome in African catfish. Our findings confirm that the African catfish stocks analyzed by us possess XX/XY sex determination mechanism only and early heat treatments with the parameters described earlier failed to affect their sex ratios.

## 4. Acknowledgements

The authors acknowledge the help of the following farms, who provided access to their stock: The Győri Előre Fisheries Co-operative (Kisbajcs, Hungary), the V95 Ltd (Nagyatád, Hungary), the Jászkiséri Halas Ltd. (Jászkisér, Hungary), and the Szarvas-Fish Ltd. (Szarvas, Hungary).

## 9. Supplementary Table

**Supplementary Table S1:** Primers used to test the applicability of recently published ‘moderately sex-associated loci’ for molecular sexing in African catfish

## 10. Supplementary Figure Legends

**Supplementary Figure S1**: Identification of the phenotypic sex by aceto-indigo carmine squash method under a stereomicroscope (Leica M205FA) with 40-100X magnification (A: female gonad, B: male gonad)

**Supplementary Figure S2**: PCR products of two putative ‘moderately male-linked loci’ (*dtna, add3*) and two putative ‘moderately female-linked loci’ (*dctn4, pcdh2ab3*) did not show any sign of sex-specific amplification profile. Molecular weight marker: GeneRuler DNA Ladder Mix (Thermo Scientific, 100 bp – 10 kb). Left – eight females, right – eight males. PCR products were separated on a 1.5% agarose gel in 1x TBE buffer.

